## Supplementary Figures for "*Myoparr*-associated and -independent multiple roles of heterogeneous nuclear ribonucleoprotein K during skeletal muscle cell differentiation"

### Supplementary Figure 1

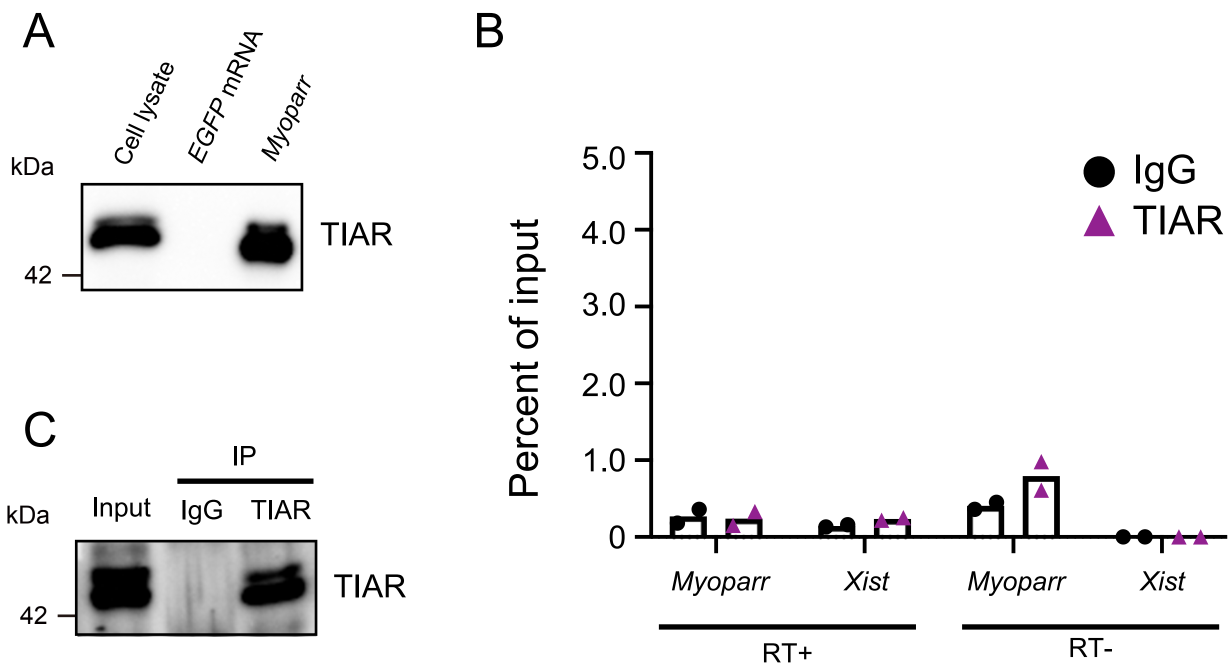

### Supplementary Figure 2

A

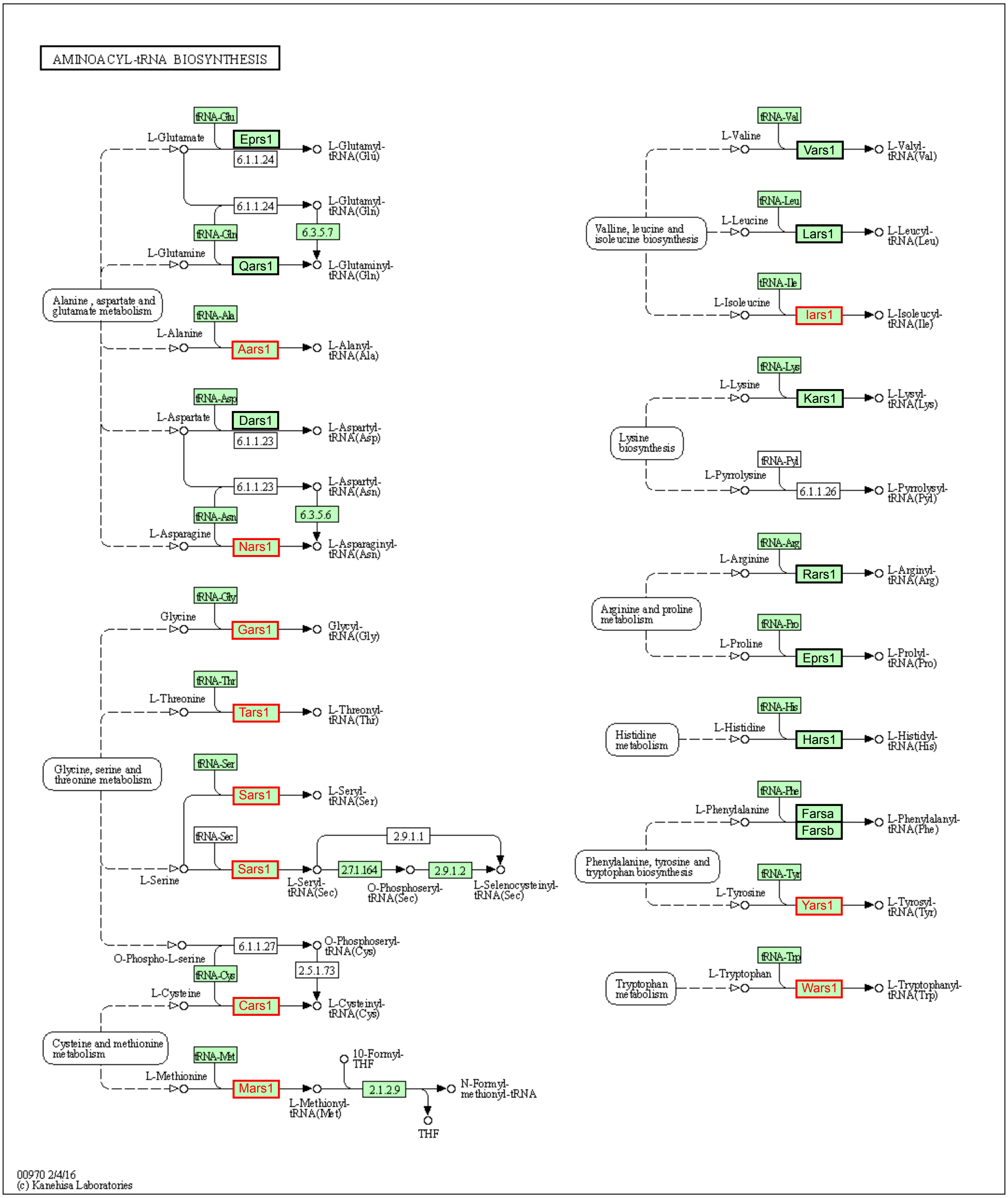


#### B Cytosolic aminoacyl-tRNA synthetases

| Gene Name | Description | log2 Foldchange | padj |
|---|---|---|---|
| *Aars (Aars1)* | alanyl-tRNA synthetase | 1.189688646 | 6.64E-11 |
| *Rars (Rars1)* | arginyl-tRNA synthetase | 0.717750186 | 0.000603064 |
| *Nars (Nars1)* | asparaginyl-tRNA synthetase | 1.037402961 | 1.73E-09 |
| *Dars (Dars1)* | aspartyl-tRNA synthetase | 0.298274648 | 0.325335115 |
| *Cars (Cars1)* | cysteinyl-tRNA synthetase | 0.872794782 | 2.21E-05 |
| *Qars (Qars1)* | glutaminyl-tRNA synthetase | 0.081217744 | 0.864024052 |
| *Eprs (Eprs1)* | glutamyl-prolyl-tRNA synthetase | 0.585228462 | 0.018216177 |
| *Gars (Gars1)* | glycyl-tRNA synthetase | 1.205275471 | 3.91E-13 |
| *Hars (Hars1)* | histidyl-tRNA synthetase | 0.413615028 | 0.200347566 |
| *Iars (Iars1)* | isoleucine-tRNA synthetase | 1.022779598 | 8.54E-07 |
| *Lars (Lars1)* | leucyl-tRNA synthetase | 0.709821097 | 0.000114578 |
| *Kars (Kars1)* | lysyl-tRNA synthetase | 0.349181856 | 0.168606343 |
| *Mars (Mars1)* | methionine-tRNA synthetase 1 | 0.900197979 | 1.01E-05 |
| *Farsa* | phenylalanyl-tRNA synthetase, alpha subunit | 0.200736108 | 0.620163384 |
| *Farsb* | phenylalanyl-tRNA synthetase, beta subunit | 0.710725176 | 0.000803764 |
| *Sars (Sars1)* | seryl-aminoacyl-tRNA synthetase | 1.078993838 | 2.66E-06 |
| *Tars (Tars1)* | threonyl-tRNA synthetase | 0.905000211 | 8.27E-07 |
| *Wars (Wars1)* | tryptophanyl-tRNA synthetase | 0.878940288 | 5.01E-06 |
| *Yars (Yars1)* | tyrosyl-tRNA synthetase | 0.772263805 | 0.000703005 |
| *Vars (Vars1)* | valyl-tRNA synthetase | 0.593311919 | 0.003371758 |

#### C Mitochondrial aminoacyl-tRNA synthetases

| Gene Name | Description | log2 Foldchange | padj |
|---|---|---|---|
| *Aars2* | alanyl-tRNA synthetase 2, mitochondrial | -0.781345711 | 0.012684624 |
| *Rars2* | arginyl-tRNA synthetase 2, mitochondrial | 0.020716521 | 0.976825609 |
| *Nars2* | asparaginyl-tRNA synthetase 2 (mitochondrial)(putative) | 0.356286565 | 0.378947013 |
| *Dars2* | aspartyl-tRNA synthetase 2 (mitochondrial) | -0.118907946 | 0.852207722 |
| *Cars2* | cysteinyl-tRNA synthetase 2 (mitochondrial)(putative) | -0.419664503 | 0.30297916 |
| *Ears2* | glutamyl-tRNA synthetase 2, mitochondrial | 0.432715891 | 0.343242224 |
| *Hars2* | histidyl-tRNA synthetase 2 | -0.125988909 | 0.813766049 |
| *Iars2* | isoleucine-tRNA synthetase 2, mitochondrial | -0.111305242 | 0.796505735 |
| *Lars2* | leucyl-tRNA synthetase, mitochondrial | -0.257452115 | 0.40655708 |
| *Mars2* | methionine-tRNA synthetase 2 (mitochondrial) | -0.463741055 | 0.383638447 |
| *Fars2* | phenylalanine-tRNA synthetase 2 (mitochondrial) | 0.24527022 | 0.65503804 |
| *Pars2* | prolyl-tRNA synthetase (mitochondrial)(putative) | 0.203237774 | 0.784806846 |
| *Sars2* | seryl-aminoacyl-tRNA synthetase 2 | -0.782886579 | 0.039853007 |
| *Tars2* | threonyl-tRNA synthetase 2, mitochondrial (putative) | 0.25526333 | 0.552363803 |
| *Wars2* | tryptophanyl tRNA synthetase 2 (mitochondrial) | 0.172212967 | 0.789056335 |
| *Yars2* | tyrosyl-tRNA synthetase 2 (mitochondrial) | 0.078853278 | 0.90065489 |
| *Vars2* | valyl-tRNA synthetase 2, mitochondrial | 0.188587495 | 0.706779801 |

### Supplementary Figure 3

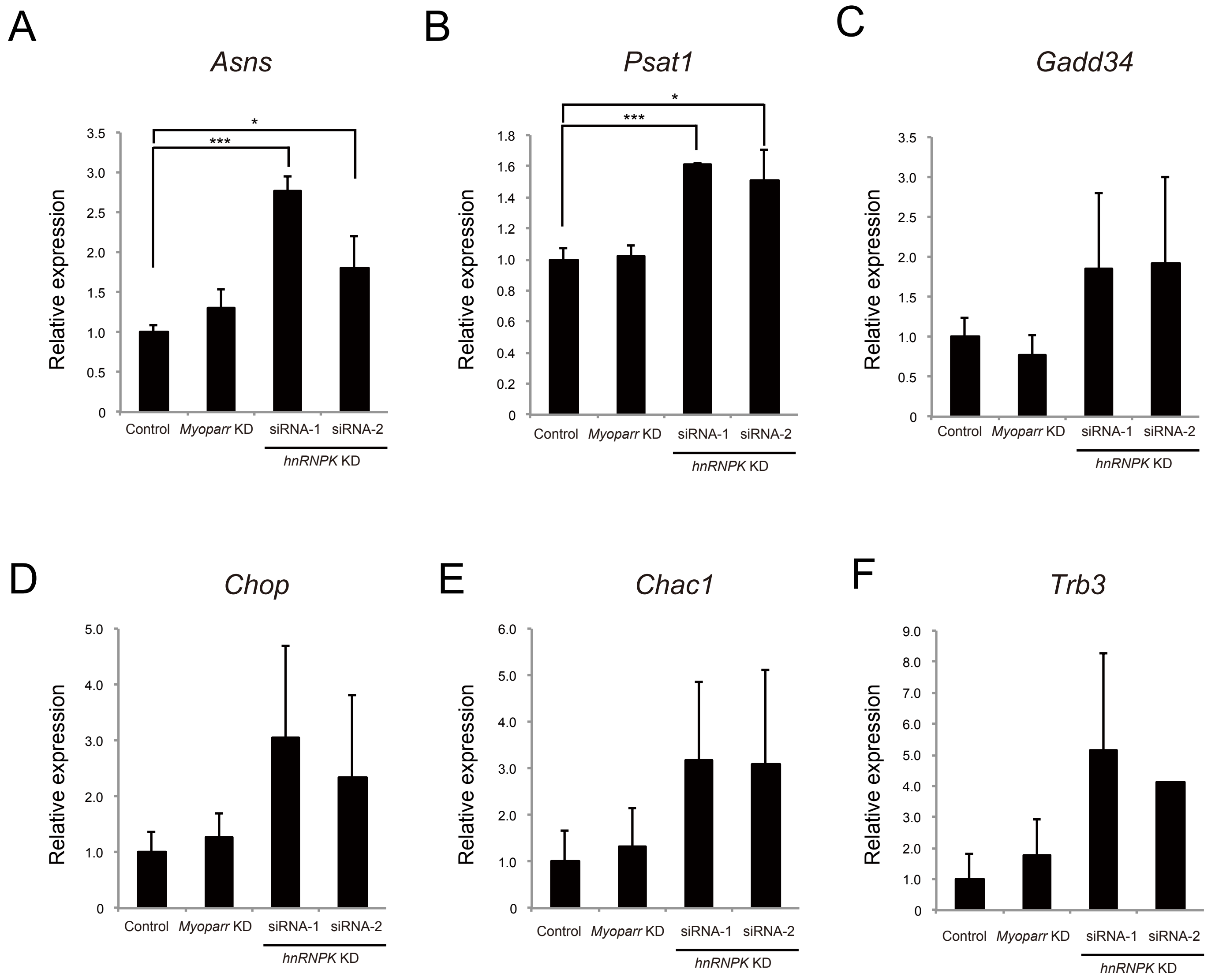

### Supplementary Figure 4

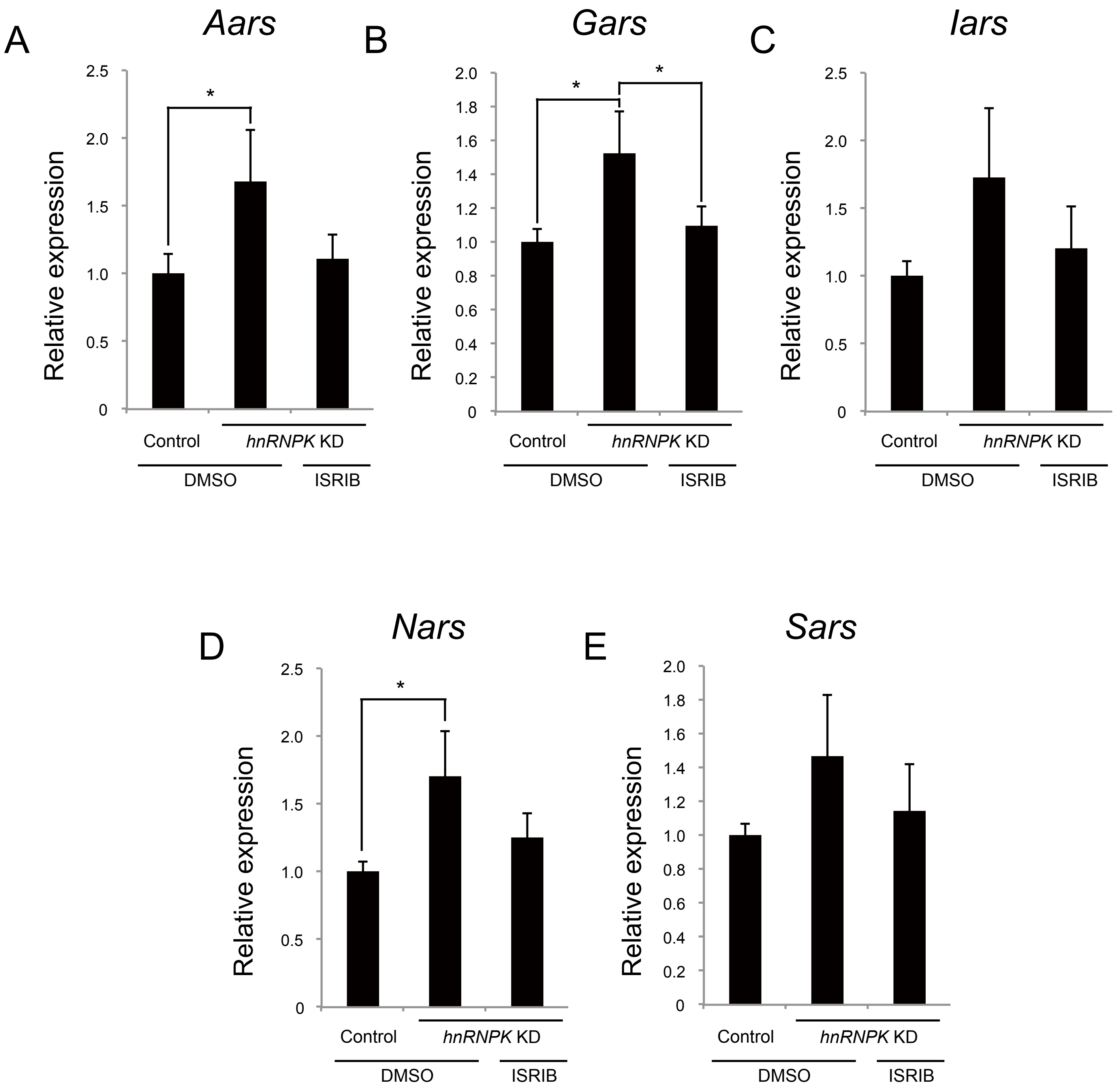

### Supplementary Table 1

| **Primers for real-time PCR analysis** | | |
|---|---|---|
| Target name | Forward | Reverse |
| *myogenin* | CAGTACATTGAGCGCCTACAG | GGACCGAACTCCAGTGCAT |
| *Myoparr* (1) | GTGCCCTATCGTCCATGGAG | CACTGACTTCACCTGACCCC |
| *Myoparr* (2) | GGGCAGGAAGGGAACAAGAA | CCCCAAATCTGGAGTGGTCC |
| *Xist* | CCCGCTGCTGAGTGTTTGAT | ATCCAGGCAATCCTTCTTCTTG |
| *Neat1* | TTGGGACAGTGGACGTGTGG | TCAAGTGCCAGCAGACAGCA |
| *Atf4* | AAGGAGGAAGACACTCCCTCT | CAGGTGGGTCATAAGGTTTGG |
| *Aars* | ATGGATGCCACTTTAACAGCA | TGGGTCGATTGTGTTTAGGAAGA |
| *Gars* | GGAGGCAGCACTTTATCCAAG | TCGGAAGCACTCTCCGTTCT |
| *Iars* | CCTCCCTTTGCTACTGGACTG | GGTGAGCGTATCTCGTAACGAT |
| *Nars* | GAGCTGTATGTATCTGACCGAGA | AAATGGTGGGAAATGGCTCTTT |
| *Sars* | CGGGTGGATAAAGGAGGGGA | TGCCCGAAATCTGCATCGTC |
| *Asns* | GCAGTGTCTGAGTGCGATGAA | TCTTATCGGCTGCATTCCAAAC |
| *Psat1* | CAGTGGAGCGCCAGAATAGAA | CCTGTGCCCCTTCAAGGAG |
| *Gadd34* | GAGGGACGCCCACAACTTC | TTACCAGAGACAGGGGTAGGT |
| *Chop* | CTGGAAGCCTGGTATGAGGAT | CAGGGTCAAGAGTAGTGAAGGT |
| *Chac1* | CTGTGGATTTTCGGGTACGG | CCCCTATGGAAGGTGTCTCC |
| *Trb3* | GCAAAGCGGCTGATGTCTG | AGAGTCGTGGAATGGGTATCTG |
| *hnRNPK* | CCGTACAGACTACAATGCCAG | GCCCTCTTCCAAGGTAGGGAT |
| *Rpl26* | GGTCTATGCCCATTCGGAAGG | TCGTTCGATGTAGATGACGTACT |

| **siRNA** | | |
|---|---|---|
| Target name | Target Sequence | |
| Control siRNA | - | Stealth RNAi siRNA Negative Control, Med GC (Thermo Fisher Scientific) |
| *Myoparr* siRNA | GATGGACCCTGTCTGATGCTCTTAA | Thermo Fisher Scientific |
| *hnRNPK* siRNA-1 (used for Figure 5, 6 and Supplementary Figure 3) | TCCCAAAGATTTGGCTGGATCTATT | MSS205172 (Thermo Fisher Scientific) |
| *hnRNPK* siRNA-2(used for Figure 5, Supplementary Figure 3, and Supplementary Figure 4) | GGAAGTGACTTTGATTGCGAGTTGA | MSS205173 (Thermo Fisher Scientific) |
